# The computation of directional selectivity in the *Drosophila* OFF motion pathway

**DOI:** 10.1101/721902

**Authors:** Eyal Gruntman, Sandro Romani, Michael B. Reiser

## Abstract

The direction of visual motion in *Drosophila* is computed by separate pathways for moving ON and OFF features. The 4^th^ order neurons T4 (ON) and T5 (OFF) are the first neurons in their respective pathways to extract a directionally selective response from their non-selective inputs. Recent functional studies have found a major role for local inhibition in the generation of directionally selective responses. However, T5 lacks small-field inhibitory inputs. Here we use whole-cell recordings of T5 neurons and find an asymmetric receptive field structure, with fast excitation and persistent, spatially trailing inhibition. We assayed pairwise interactions of local stimulation across the receptive field, and find no active amplification, only passive suppression. We constructed a biophysical model of T5 based on the classic Receptive Field. This model, which lacks active conductances and was tuned only to match non-moving stimuli, accurately predicts responses to complex moving stimuli.

## Introduction

Visual motion is not directly measured, rather it is computed by neuronal circuits downstream of photoreceptors. This computation is fundamental to the extraction of many visual features, is a local operation implemented by a small circuit, and is simple enough to be approximated by compact algorithms. Although several different algorithms have been proposed for generating directionally selective responses across species, they all share three common elements: (1) spatially offset inputs with a (2) temporal asymmetry between them, that are (3) non-linearly combined. These algorithms generate Directional Selectivity (DS) by enhancing responses in the preferred direction (seen in flies: (Fisher et al., 2015; Salazar-Gatzimas et al., 2016)), suppressing responses in the null, or non-preferred direction, or by a combination of both (also from flies: (Haag et al., 2016; Leong et al., 2016; Strother et al., 2017), reviewed in (Yang and Clandinin, 2018)). In *Drosophila*, local luminance increments (ON) and decrements (OFF) are processed by largely separate circuits, with motion being computed separately in each pathway (Behnia et al., 2014; Clark et al., 2011; Joesch et al., 2010; Silies et al., 2013; Strother et al., 2014; Takemura et al., 2014). The first neurons to generate directionally selective responses are the T4 cells of the ON pathway and the T5 cells of the OFF pathway (Maisak et al., 2014; Serbe et al., 2016; Strother et al., 2017) (**Fig. 1A**).

**Figure 1.**
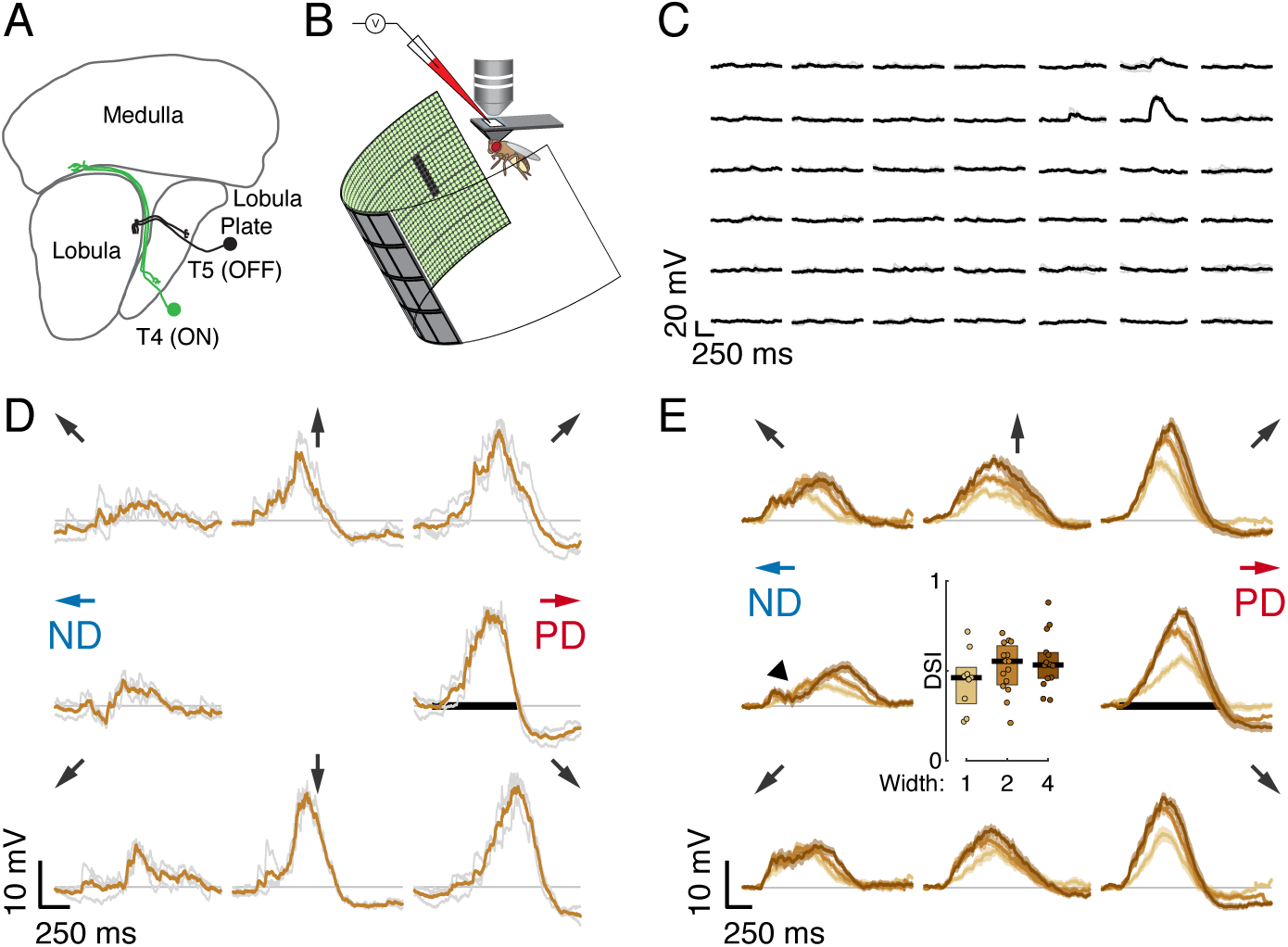
Whole-cell recordings of T5 neurons show small-field, directionally selective responses. (**A**) Schematic of the *Drosophila* visual system with an example T4 (ON) and T5 (OFF) neuron. (**B**) Schematic of experimental setup. Whole-cell recordings were targeted to soma of labeled T5 neurons. (**C**) Responses to 200ms OFF square flashes (∼11° × 11°) from an example cell. Each subplot is the response from a different location on the LED display, which subtended 216° (azimuth) × 72° (elevation) of the visual space. Individual repeats in gray (n=3 trials), mean in black. (**D**) Responses from the same cell in **C** to a 2-pixel wide dark bar (2 × 9 LEDs, ∼5° × 20°) moving in eight directions at 28°/s (80ms/pixel) through the center of the receptive field. Repeats in gray (*n* = 3 trials), mean in brown. Black horizontal bar indicates the stimulus duration. PD indicates the Preferred Direction and ND indicates the Null Direction. (**E**) Baseline-subtracted responses (*n* = 17 cells) to a moving bar of with 1, 2, and 4 pixels (2.25°, 4.5°, and 9°), aligned to the PD of each cell (mean ± SEM). Arrows represent the direction of stimulus motion. Black horizontal bar indicates stimulus presentation. Arrowhead indicates dip in ND response. Inset: DSI = (PD_max_ – ND_max_)/ PD_max_ for moving bar responses (*n* = 9, 15, 14 cells).

T4 neurons use both local excitatory and inhibitory inputs to generate directionally selective responses (Gruntman et al., 2018; Haag et al., 2017; Strother et al., 2017; Takemura et al., 2017). A recent connectomic study has characterized all the columnar inputs to T5 neurons (Shinomiya et al., 2019) and a functional imaging study revealed they all depolarize in response to OFF stimuli (Serbe et al., 2016). However, transcriptional profiling of these neuron types shows that these inputs are all cholinergic, and therefore unlikely to provide local inhibitory input (Davis et al., 2018). Does the OFF pathway use a different algorithm to compute motion? To address this question, we performed whole-cell recordings of T5 neurons while presenting visual stimulation. First, we used single bar flashes to map the first-order Receptive Field (RF). These stimuli, which do not contain any motion information, allowed us to map the spatial distribution of excitatory and inhibitory inputs and revealed that T5 neurons receive local inhibitory inputs. Next, we used many variants of pairwise bar flashes, an elementary motion stimulus, and consistently found only a single mechanism responsible for DS generation. Next, we constructed a conductance-based model for a T5 cell and used it to predict responses to an array of visual stimuli. The model is comprised of fast excitation and slow inhibition spatially offset to the trailing side of the RF (Gruntman et al., 2018). We show that our model, constructed only from the first-order RF responses that contain no motion information, predicted responses to complex dynamic stimuli, such as moving bars and drifting gratings. Finally, we analyzed the behavior of this class of computational models. We find that our predictive models of T5 neuron responses use parameters that are close to optimal for detecting the direction of motion.

## Results

### Whole-cell recordings of T5 neurons show small-field, directionally selective responses

We measured visual responses of T5 neurons by targeting in-vivo whole-cell electrophysiological recordings to their GFP-labelled somata and presenting stimuli on a hemi-cylindrical LED display (**Fig. 1B**). We confirmed the identity of the labeled neurons as T5 cells by recording reliable depolarizations in response to small OFF flashing squares (∼11° × 11°; pixels turned off from an intermediate-intensity background). These flashing squares were also used to localize the RF center (**Fig. 1C**) in a step-wise process that mapped the maximal response position for each recorded neuron by probing smaller areas at higher resolution until the peak response was localized to a single pixel (∼2° × 2°) of the display. Having localized the RF center, we then evaluated DS for each neuron by measuring responses to bars of 3 widths moving through the RF center along 8 directions. The bars moved at the speed (28°/s) that produced the largest directionally selective responses in T4 neurons (Gruntman et al., 2018). Each of the 17 T5 neurons we recorded showed clear preferred and null directions (PD and ND) of motion, with relatively wide tuning (i.e. similar magnitude responses in the directions ±45° away from the PD; individual neuron example **Fig. 1D**; population responses **Fig. 1E**). The PD response shows a clear hyperpolarization following the depolarizing peak, and the ND response shows a dip preceding the depolarization peak (arrowhead in **Fig. 1E**). We quantified these responses with a Directional Selectivity Index (DSI = [PD_max_ – ND_max_]/ PD_max_), and found that this measure is significantly different from zero for the 3 widths tested (**Fig. 1E**). We note that the smallest bar presented (1-pixel wide, ∼2°) generated a clear directionally selective response, but did not induce the prominent hyperpolarization observed for the wider bars. We further note that although wider bars evoked stronger responses, they did not increase the DSI (**Fig. 1E**), because these stronger stimuli also evoked a corresponding increase in the ND response magnitude. The response dynamics and the width of the directional tuning are similar to our T4 responses to moving bars (Figure 1 of Gruntman et al. (2018)).

### T5 receptive field is comprised of spatially offset excitatory and inhibitory inputs

Classic studies of mammalian directionally selective neurons used decomposable motion stimuli to map spatial responses to the individual components (the “first-order RF”), and to their pairwise interactions (the “second-order RF”). This procedure was important in ruling out competing models for generating DS (Emerson et al., 1992; 1987; Jagadeesh et al., 1993). In our previous work (Gruntman et al., 2018), we adapted this approach to T4 neurons, and found that a fine-scale characterization of the neuron’s first-order RF was incompatible with the Hassenstein-Reichardt model (Hassenstein and Reichardt, 1956), the predominant model for computing DS in insects. We wondered whether the differences between the signs of T4’s and T5’s columnar inputs would also produce functional changes in the first-order RF. We therefore used on-line stimulus generation to map the spatial and temporal properties of T5 receptive fields using flashing bars presented along the identified PD-ND axis of each cell (**Fig. 1**). We used bars of width 1, 2, and 4 pixels (corresponding to 2.25°, 4.5°, and 9° of visual angle), and our RF maps are plotted (**Fig. 2**) along a stimulus position axis, where a unit change in position is equivalent to a 1-pixel movement of a width 1 bar. Although these single position bar flashes lack directional information, the temporal and spatial structure of T5 responses to them make up the first-order RF that is used for all further comparisons, since more complex stimuli can be comprised of different concatenations of single position flashes.

**Figure 2.**
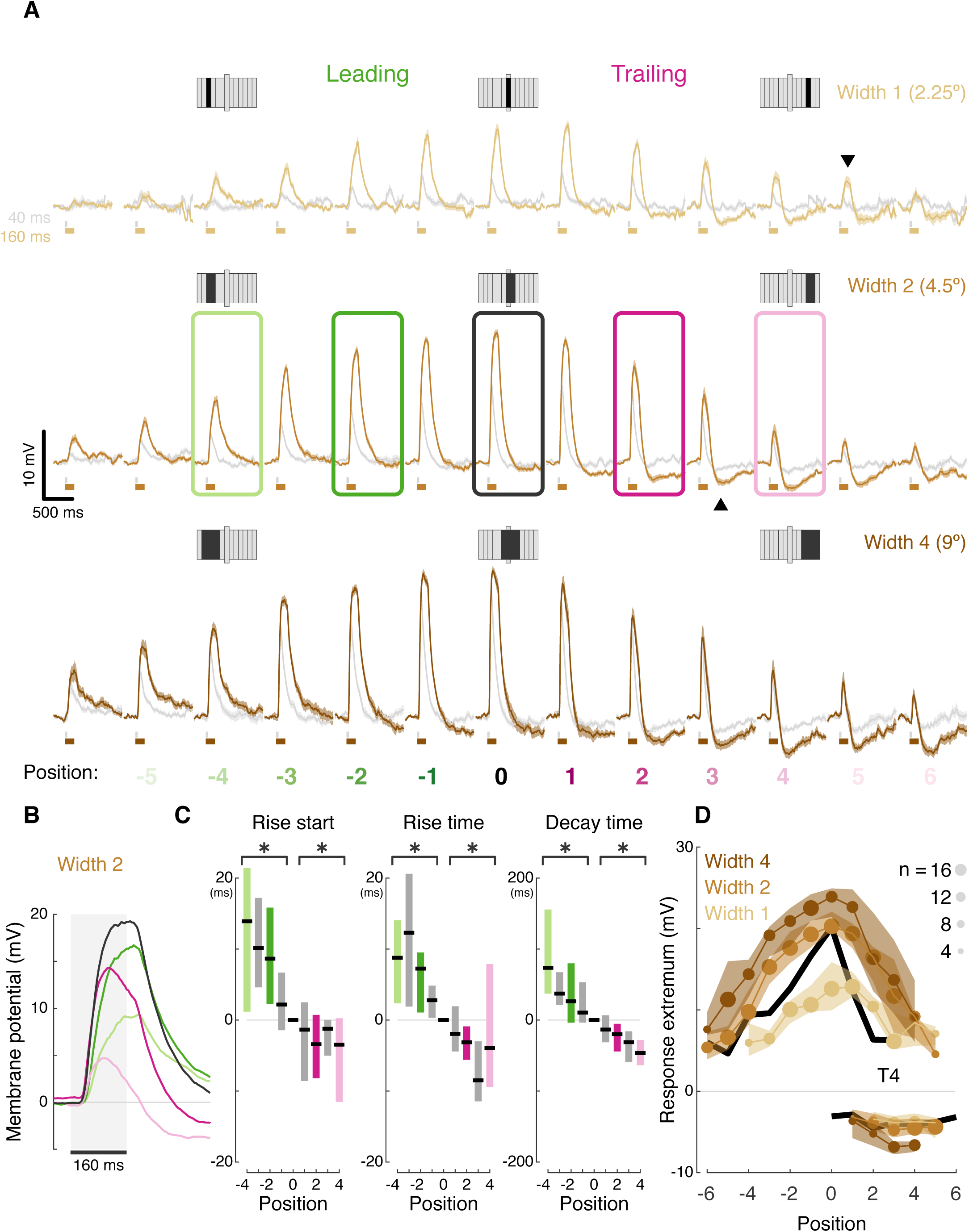
T5 receptive field is comprised of spatially offset excitatory and inhibitory inputs. (**A**) Averaged, baseline-subtracted responses (mean ± SEM) to bar flash stimulus at the indicated positions (numbered below, examples schematized above) along the PD–ND axis of each cell (*n* = 17 cells) aligned to the center excitatory position (=0, see Methods for details). Responses to 40ms flashes in gray; responses to 160ms flashes are colored. Enlarged bar in each stimulus schematic marks center position. Downwards arrowhead: example non-linearity in responses to stimuli of different durations. Upwards arrowhead: example non-linearity in hyperpolarization in response to stimuli of different durations. (**B**) Mean responses from indicated positions in **A** of width 2 bar flashes, aligned to stimulus presentation (gray rectangle). (**C**) Response rise start time (time to reach 10% of max), rise time (10%-50%), and decay time (80%-20%) for positions surrounding the receptive field center (calculated for 160ms flashes of width 2 bars). Results presented as differences from the central position (* indicates significantly above/below zero for pooled positions from the leading/trailing side respectively, one-sided unpaired t-test, p < 0.001; n=17 cells). (**D**) Maximum depolarizing and hyperpolarizing responses at each stimulus position for 160ms bar flashes of all 3 widths. Dots correspond to median response (size indicates the number of cells for each position), shaded regions demarcate upper and lower quartiles. Superimposed black line represents the median T4 results for width 1 ON bars (Gruntman, et al. 2018).

Aligning the responses from all the cells based on the position of the peak depolarization (**Fig. 2A**) reveals T5’s (first-order) RF structure: (1) Excitatory responses dominate the center, with increased response magnitude for stronger stimuli (wider bars or flashes of longer duration). This increase is not linear. For example, at certain positions responses cannot be detected for weak stimuli but clear responses can be seen for stronger stimuli (e.g. compare bar width 1 at position 5, gray vs. brown traces, downwards arrowhead). (2) Inputs along the PD-ND axis are spatially asymmetric around the RF center. On the leading side of the RF the responses reflect excitatory input (**Fig. 2A**, green positions), while on the trailing side, responses show a mixture of excitation and inhibition (**Fig. 2A**, pink positions). This general structure is similar to T5 measurements in a recent paper using voltage imaging (Wienecke et al., 2018). Although stronger stimuli (wider bars or longer duration flashes) induce a stronger hyperpolarizing component in the response, this effect is also non-linear. For example, average traces in position 3 (bars of width 2) show no hyperpolarization for a 40ms flash (gray trace), but a prominent hyperpolarization when the flash duration is 160ms (brown trace, **Fig. 2A**, upwards arrowhead).

The unipolar structure of *Drosophila* neurons precludes the use of voltage clamp to separate the inhibitory and excitatory conductances in the responses. However, a closer examination of the response dynamics from the leading and trailing side of the RF reveals a temporal interaction between excitation and inhibition. Long duration responses on the leading side show a plateau-like depolarization peak, while responses on the trailing side show a peak that rapidly decays (**Fig. 2B**, pink vs. green responses). These decaying response peaks likely arise from a ‘competition’ between fast-rising, fast-decaying excitatory inputs and slow-rising, slow-decaying inhibitory inputs. At stimulus onset, the fast excitation is dominant, which induces the depolarization peak. As the slower inhibition increases with time, so does the depolarization decay. At stimulus offset, excitation decays rapidly, while inhibition persists—as is evident in the post-stimulus hyperpolarization (**Fig. 2B**).

Spatially asymmetric inhibition also induces faster decaying responses on the RF’s trailing side (**Fig. 2B** pink vs. green, quantified in **Fig. 2C**). In T4 neurons, we showed that the asymmetric, slow-decaying inhibition is the main contributor to DS generation, since ND movement first activates the faster-decaying depolarizing response components, which in turn lead to less efficient summation and smaller net depolarization (Gruntman et al., 2018). Therefore, the faster decay times on the trailing side of T5 neurons were also the likely result of inhibition temporally sharpening responses. Unlike in T4, trailing-side responses started sooner and rose faster than leading-side responses (**Fig. 2C**; single sided t-tests, p < 0.001 for all conditions). It is unclear whether these differences in rise start and rise time play a role in generating directionally selective responses, or whether they simply reflect a wider range of spatially offset excitatory input types in T5 cells (Shinomiya et al., 2019).

Plotting the peak depolarization and hyperpolarization for each cell at each position for long duration flashes of all widths illustrates the RF structure described above: excitatory component decaying symmetrically from the center, and an asymmetric inhibitory component on the trailing side (**Fig. 2D**). This structure is comparable to the RF structure we measured for T4 cells (**Fig. 2D**, black curves). Note that the T4 RF was mapped using responses to a bar of width 1, suggesting a potential size-sensitivity difference between the ON and the OFF pathways (consistent with Haag et. al. (2017)). The high degree of similarity in the first-order RF between T4 and T5 cells suggests that both cell types receive similar (functional) inputs. But do they use a similar DS mechanism? To test this directly we next characterized T5 responses to elementary motion stimuli.

### T5 neurons generate directional selectivity using only ND suppression

Sequential presentation of two adjacent bar flashes, sometimes referred to as two-step apparent motion, is a common stimulus used to map the second-order RF of directionally selective cells by comparing the superposition (summation shifted in time) of the first-order responses with the responses to the sequential, two position stimulation (**Fig. 3A**). This comparison is a direct test for the relative contribution of either PD enhancement or ND suppression in generating the directionally selective response (**Fig. 3A**). To illustrate the general response properties, we focus on responses to width 2 bars that were each presented for 160ms (**Fig. 3B,C**). Responses on the leading side of the RF, up to and including the center (position 0), did not exhibit DS. Since both PD and ND combinations evoke a similar maximal response (e.g. -3 to -1 versus -1 to -3, horizontal lines) the neurons are effectively ‘motion blind’ in this region of the RF. Responses to combinations that included the trailing side of the RF, where inhibition was detected (**Fig. 2D**), exhibited a larger maximal response for the PD combination, and therefore exhibited DS (e.g. **Fig. 3B** 1 to 3 in red versus 3 to 1 in blue, horizontal lines).

**Figure 3.**
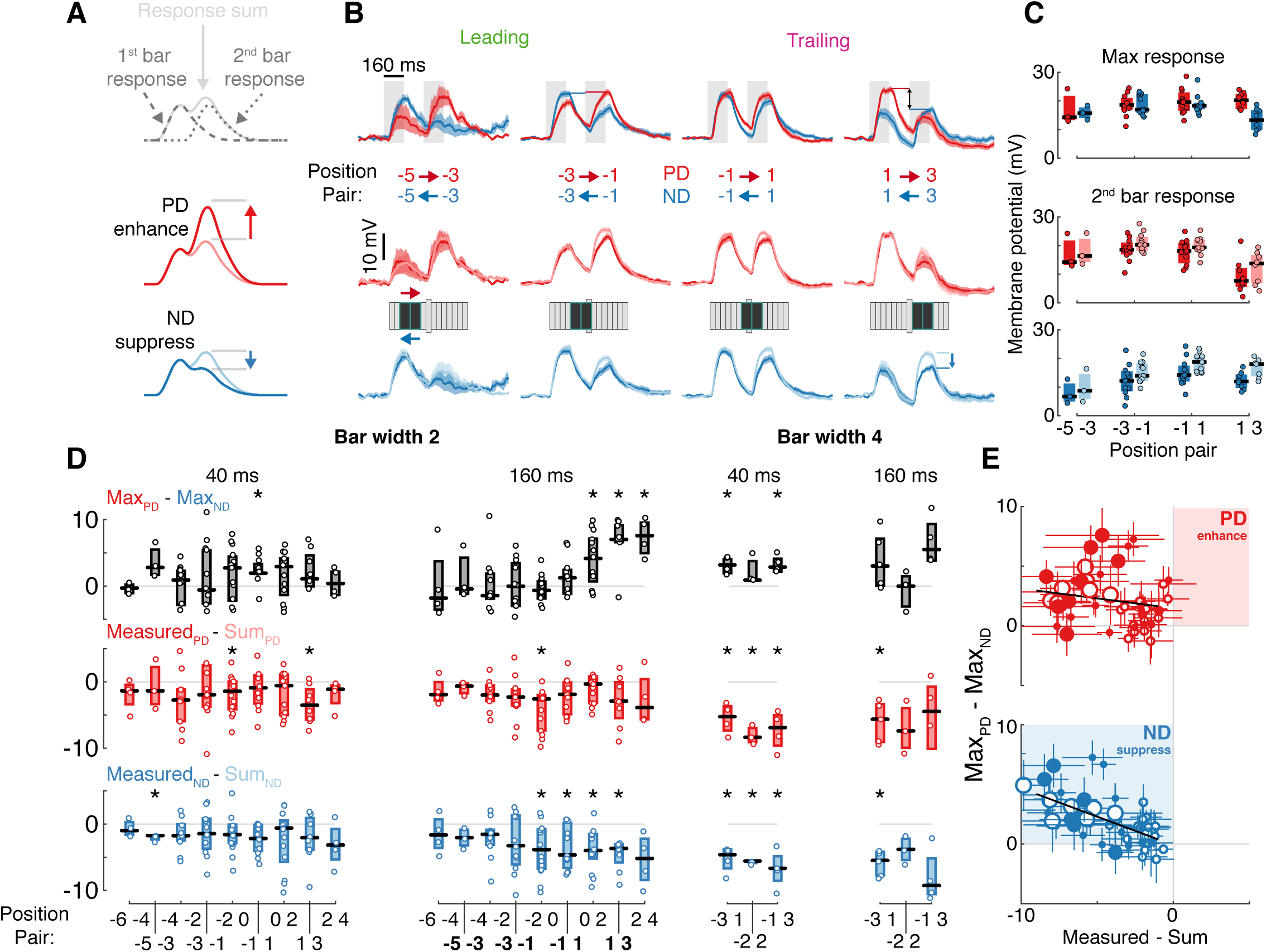
T5 neurons generate directional selectivity using only ND suppression. (**A**) Schematized responses to the elementary motion stimulus of sequential bar pair flashes. Response could be the sum of the responses to the individual flashes (top), could show preferred direction, PD, enhancement (middle) or null direction, ND, suppression (bottom). (**B**) Baseline-subtracted responses (mean ± SEM) to bar pair combinations presented at different positions along the PD–ND axis (*n* = 3,11,11,8 cells). Top: responses to PD and ND bar pairs. Middle: responses to PD bar pairs (same as top) with the temporally aligned sum of the responses to the component bar flashes (pink). Bottom: responses to ND bar pairs (same as top), sum of responses to component bar flashes in cyan. Stimulus presentation interval indicated with the gray rectangles. Stimulus schematic shows positions of both bars (0 position depicted by elongated bar). (**C**) Boxplot summary of response maxima for **B** (same color conventions). Top: response maxima, Middle and Bottom: second bar maxima for measured and summed responses. (**D**) Boxplot summary of response maxima differences for all (non-overlapping) bar pair stimuli presented. Top: difference between PD and ND, positive value indicates a directionally selective response. Middle: difference between second bar response maxima for measured and summed PD responses. Positive value indicates PD enhancement. Bottom: same as middle, but for ND. Negative value indicates ND suppression (* indicates mean significantly differs from zero, unpaired t-test corrected for multiple comparisons by controlling for the false discovery rate with q = 0.075). Boldface positions are presented in **B** and **C.** (**E**) Comparison of directional selectivity versus linearity of response for all presented bar pair combinations, including data from **D** and responses to overlapping positions (see Methods). Each dot corresponds to the mean (of n ≥ 3 cells) response differences for each position pair (± SEM). Marker size indicates bar width (small for 2, large for 4-pixel wide), marker fill indicates duration (empty 40ms, filled 160ms). Results of linear regression in black (non-significant slope for PD_measured-sum_ vs. Max_PD-ND_, [-0.402, 0.073], 95% confidence interval; significant slope for ND_measured-sum_ vs. Max_PD-ND_, [-0.641, -0.301], 95% confidence interval). See also Fig. 3 Fig. Supp. 1 for more details of bar width 4 responses.

To further explore the mechanism underlying DS we compare each 2-bar stimulus response (**Fig. 3B** middle and bottom, darker traces) to the superposition of the component responses (**Fig. 3B** middle and bottom, lighter traces). In the leading side of the RF, where T5 is ‘motion blind,’ we find that the sequential responses are well approximated by the sum of the individual responses. However, in the trailing side of the RF, where T5 generates directionally selective responses, this comparison reveals suppression of ND responses (blue arrow), as can be seen in the comparison between responses to the second bar (whose location provides the directional component of the stimulus pair; **Fig. 3C**). We note that there are no conditions where the summed responses are smaller than the measured responses. In other words, no combinations of 4.5° bars that showed PD enhancement. A recent paper using calcium imaging reported PD enhancement in both T4 and T5 cells, but only for stimuli above a certain size (> 6° for T5s; (Haag et al., 2017)). However, our findings hold for all of the apparent motion conditions we tested: fast (40ms) and slow (160ms), bars of width 2 (4.5°) and width 4 (9°), and sequential positions that were either adjacent or overlapping (**Fig. 3D,E** and **Fig. 3 - Fig. Supp. 1**). Although PD responses exhibited suppression (**Fig. 3D**, middle), these stimuli still evoked directionally selective responses in trailing side positions due to the asymmetric structure of the RF and the even larger suppression of ND stimulation (**Fig. 3D**, top).

To summarize the responses to all of these different pairwise combinations we plot the difference between the peak responses of each PD and ND sequence pair (a measure of DS, with a positive value indicating PD preference) against the difference between the measured response and the sum of the component responses (indicating suppression when negative, enhancement when positive). Although we used an expansive stimulus set, we found numerous combinations showing ND suppression and not a single condition showing an enhanced PD response. Furthermore, we found a significant correlation between the magnitude of ND suppression and the magnitude of the directionally selective response (more suppression correlates with more DS), while no such correlation was found for PD responses (**Fig. 3E**; ND_Measured-Sum_ vs. Max_PD-ND_ R2: 0.396, slope 95% confidence interval: [-0.64, -0.31]; PD_Measured-Sum_ vs. Max_PD-ND_ R2: 0.02, slope 95% confidence interval: [-0.402, 0.073]). Taken together, these data provide strong evidence for the suppression of ND motion as the only mechanism through which DS is generated in T5 neurons.

### A conductance-based model quantitatively predicts directionally selective responses

We constructed a conductance-based neuronal model to test our intuitive proposal regarding the generation of directionally selective responses in T5 cells. The model includes an excitatory and an inhibitory conductance that are combined using a biophysically inspired non-linearity (**Fig. 4A**). The model parameters primarily define the spatial and temporal filters for each conductance and the membrane’s time constant (**Table S1** and Methods). Based on our previous modeling of T4 responses (Gruntman et al., 2018), we hypothesized that the passive integration of these conductances would be sufficient to explain T5’s response dynamics. Our objective in constructing the model was not only to generate DS responses, but to explore whether the information contained in the responses to non-moving stimuli is sufficient to predict the neuron’s responses to more complex stimuli, such as drifting gratings.

**Figure 4.**
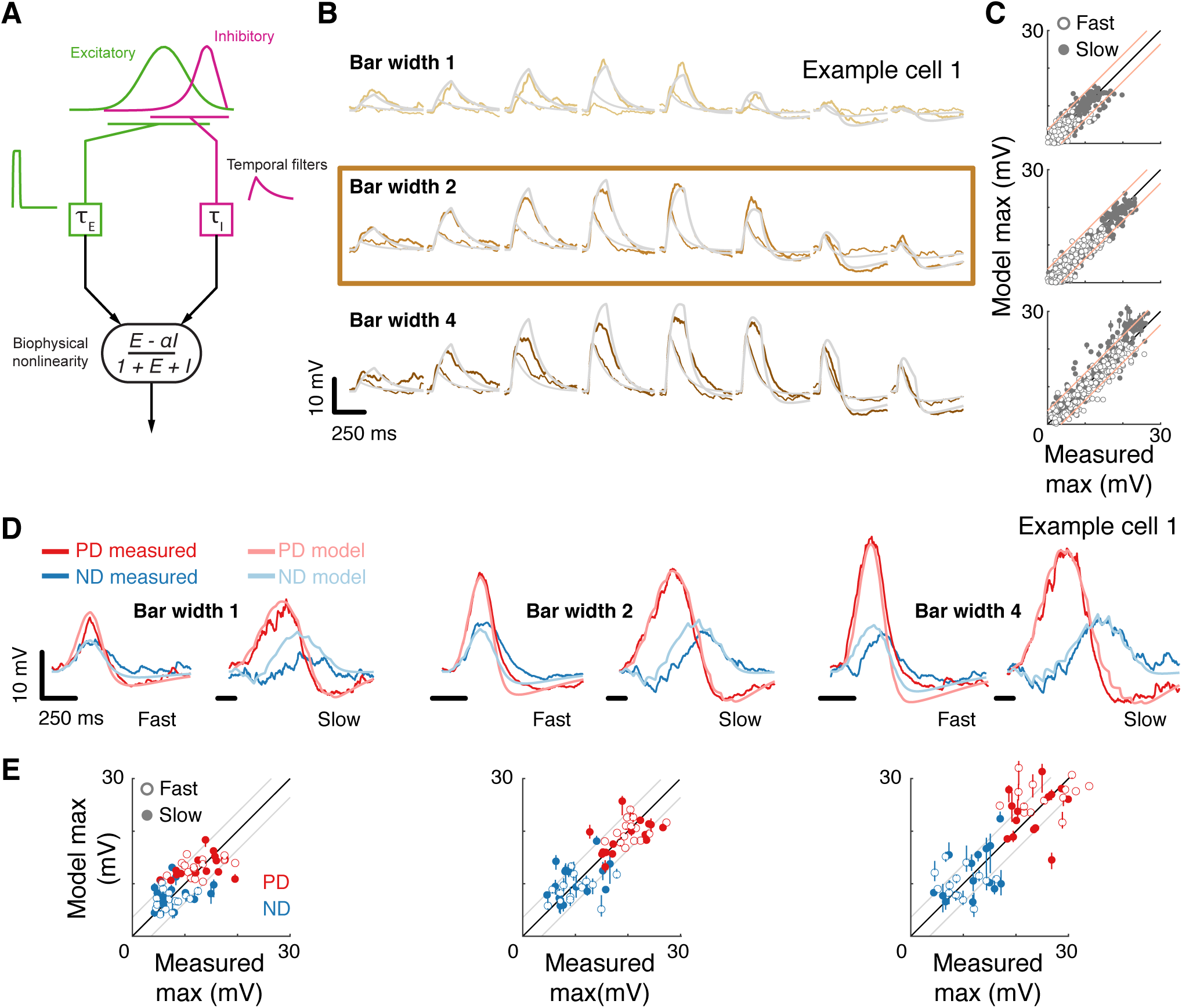
A conductance-based model quantitatively predicts directionally selective responses. (**A**) T5 Model schematic with fast spatially symmetric excitation and slow, trailing-side asymmetric inhibition. (B) Mean measured responses to single bar flashes of 3 widths and 2 flash durations at 8 different positions (in colors) compared to predicted model responses (gray) from an example cell (same as Fig. 1). Model parameters were fit to each cell using only responses from bars of width 2 (brown frame). (**C**) Peak measured response compared to the peak of the predicted response for all bar flash stimuli from all cells and positions (empty marker denotes short duration flash and filled denotes long flashes). Each dot represents the mean peak response (for top 10 of 1000 optimization solutions for each cell, estimated by fit error), while vertical lines (most obscured by markers) represent maximal and minimal values of the peak responses. Diagonal lines surrounding the unity line denote +/-the upper quartile of the Mean Absolute Deviation (MAD) of responses to repeated presentation of the same stimuli (see Methods). (**D**) Mean measured responses to moving bars of 3 widths and 2 speeds overlaid with predicted responses for same example cell and the same model parameters as in **B**. (**E**) Peak measured responses compared to the peak of the model’s predicted responses for all moving bars (grouped by bar width). Plotting conventions as in C; except here PD in red, ND in blue. See also Fig. 4 - Fig. Supp. 1 for distributions of individual cell measurement and model prediction responses.

We fit model parameters using an iterative, non-linear, least squares optimization procedure to minimize the difference between the numerical simulation and measured responses for width 2 bar flashes (**Fig. 4B**). Importantly, since we fit only based on responses to static flashing bars, the model parameters are not influenced by any motion-related responses. The model’s responses to any stimulus are then simply the result of passive integration of excitatory and inhibitory conductances injected with a temporal pattern determined by the spatial and temporal structure of the stimulus (see Methods). Since our aim was to predict responses to more complex stimuli and since these are sensitive to stimulus position within the RF, we chose to fit model parameters using responses from individual neurons, and not a population average. As a consequence of this approach, we made two modifications to our previous model: (1) we use an asymmetric spatial profile for inhibition and (2) we fit the reversal potential for inhibition as an additional parameter. The former change added flexibility to the spatial profile of inhibitory inputs, which was unnecessary when fitting the population average; the latter change allowed for a non-spatial adjustment to the relationship between excitation and inhibition.

The size of the stimuli used to probe directionally selective cells is a prominent confound when comparing results from different studies, which has resulted in conflicting interpretations about the possible algorithms implementing DS. For example, a previous study using Calcium imaging only observed PD enhancement in T5 cells for stimuli above 6° (Haag et al., 2017). Since we fit our parameters using responses to width 2 bar flashes alone, we postulated that if increasing the width of the bar above a certain size evoked qualitatively different responses, we would see a clear discrepancy between predictions and recorded responses for width 4 bars, but not for width 1 bars. **Fig. 4B** shows the results from an example cell, with the recorded responses and model predication overlaid for 3 bar widths and with both fast and slow flash durations. The model predicts the magnitude and dynamics of the responses to all 3 widths, with no systematic difference for the larger or smaller width bars. The broad accuracy of these predictions can be seen when comparing maximal (peak) responses between model and measurements across all the cells (**Fig. 4C**). The accuracy of model predictions to measured data will, in part, be limited by trial-to-trial variability in the recorded responses to identical stimuli. We have used this variability as a simple bound on the accuracy of model predictions, by plotting the +/-upper quartile of the mean absolute deviation (MAD) across cells on either side of the diagonal (identity) line (**Fig. 4C**). Since width 2 bars were used for fitting model parameters, the spread around the diagonal is narrowest for this condition, with most simulation results falling within the ‘MAD bounds’ (see Methods and **Fig. 4 – Fig. Supp. 1**). However, the spread around the diagonal for width 1 or width 4 is only slightly broader, confirming that the responses across these bar widths can be predicted without further modifications to the model.

Our simple model accounts for responses to local flash stimuli of multiple widths (containing no directional information), but how well does it predict responses to moving bars of different widths? The example traces in **Fig. 4D** are from the same cell as in **Fig. 4B**, now simulated for moving bar stimuli (same model parameters as above). We note that for these stimuli, the model predicts not only the magnitude of the maximal response in both directions, but also captures the dynamic structure of the response traces. For example, the simulated results to slow ND motion all show a hyperpolarizing dip before the depolarizing peak, but this dip is absent from ND motion traces for fast motion. This is explained by the slow inhibitory conductance, which cannot contribute much to the early component of the response to fast motion. To test our model’s predictions, we again compared the maximal responses between model predictions and measured data. Notably, if PD enhancement contributed to generating DS responses to moving bars of width 4 in our measurements, we would see a consistent underestimate of PD motion by the simulation. However, the predicted responses for both PD and ND motion show the same symmetrical spread around the diagonal (**Fig. 4E**, right), again confirming that a single, simple mechanism can account for DS responses to bars of different widths.

### Conductance-based model recapitulates responses to complex spatial and temporal stimuli

Given the strong correspondence between model predictions and response measurements for bar flashes and moving bars across speeds and widths, we challenged our model with more complex stimuli. Bar flashes have a simple spatial and temporal structure. Moving bars are a more complex stimulus, but they are still comprised of sequential activation of adjacent positions. The stimulus presented in the example in **Fig. 5A**, flashes of gratings in different phases, has a non-contiguous spatial structure, requiring spatial integration from different RF regions. As can be seen from the example cell (same cell and parameters as in **Fig. 4**), our model also accurately predicted responses to these grating flashes. Importantly, when the visual input stimulated both the excitatory and the inhibitory fields (E or I above stimulus schematic in **Fig. 5A**), the model predicted the dominant conductance, both in this individual example and in the population (**Fig. 5B**).

**Figure 5.**
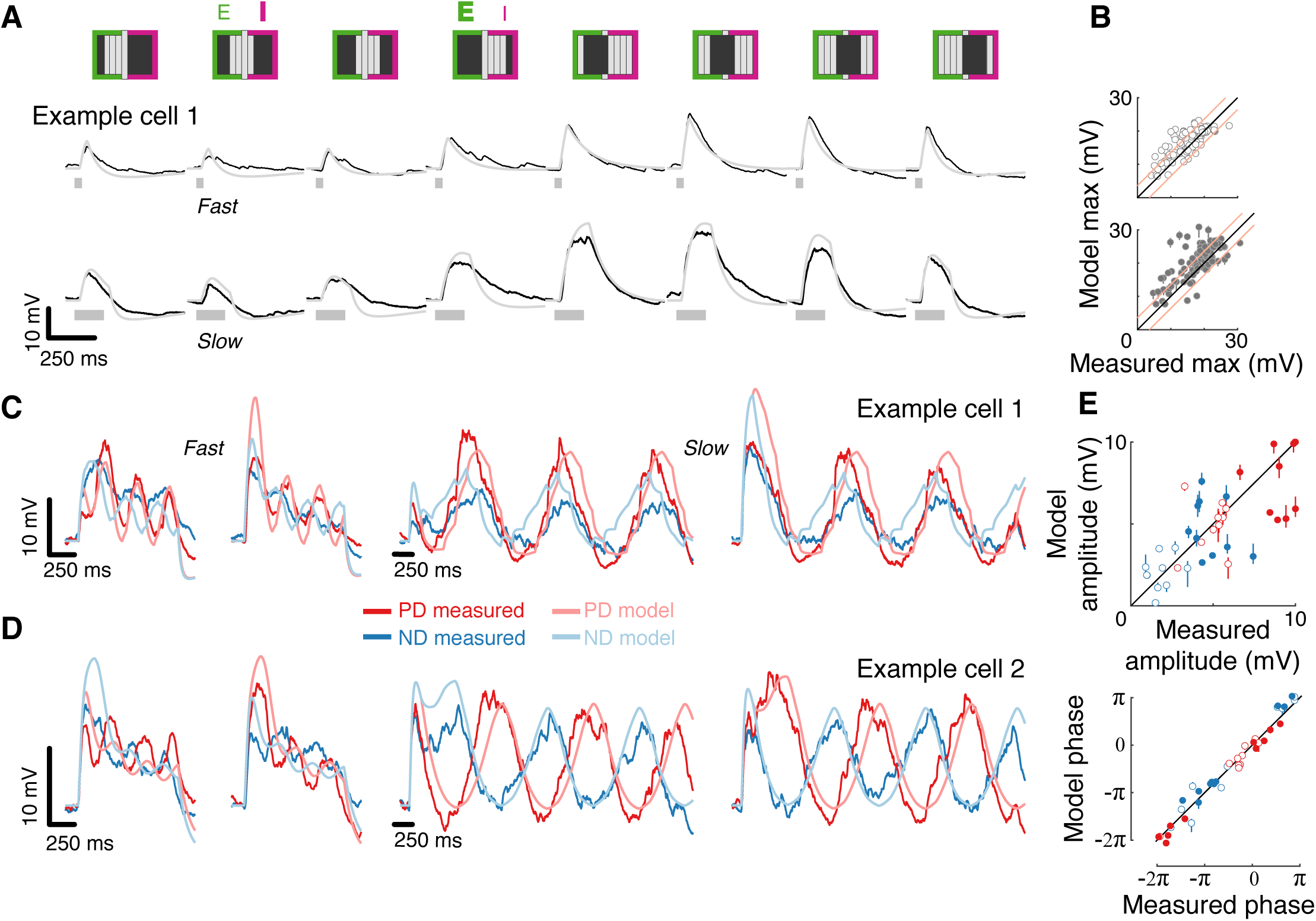
Conductance-based model recapitulates responses to complex spatial and temporal stimuli. (**A**) Mean measured responses to fast (40ms) and slow (160ms) flashes of grating stimuli (dark and background brightness level) in different phases compared with model predictions (same example cell, same model parameters as in **Fig. 4B)**. Stimulus schematic above traces. In 2 phases, the expected relative contribution (bold > normal) of excitation and inhibition based on the position of the dark bars is denoted. (**B**) Peak measured response compared to the peak of the model predicted response, for flashes of all grating phases, grouped by flash duration. Plotting conventions as in **4C** (only results from cells with a non-diagonal PD-ND axis are shown; n=5, see Methods). (**C**) Mean measured responses to grating stimuli moving at 2 speeds (temporal frequency of 3.125Hz and 0.78Hz, 40ms and 160ms steps), with 2 different starting phases, compared to model prediction for the same example cell and model parameters as **4B**. (**D**) As in **C**, only from a different example cell. Note the difference in phase relations between PD and ND responses for cells in **C** and **D**. (**E**) Top: Fit amplitude values for measured response compared to the model predicted responses for all grating stimuli (see Methods for details). Plotting conventions as in **4E**. Bottom: fit phase values for measured responses compared to model prediction responses.

Next, we challenged our model with drifting gratings, the most complex stimulus we presented. This stimulus has a non-contiguous spatial structure that is swept across the RF in a temporally cyclic manner. We simulated responses to square wave grating (composed of dark and background level bars) moving at 2 different speeds and starting from 2 different phases (**Fig. 5C**). Although the model parameters were fit using only the static stimuli responses (**Fig. 4B**, middle), the model’s predictions capture several aspects of the recorded responses: the transient responses to the appearance of the grating (that differ dramatically between the two phases), the amplitude of the oscillation (compared in **Fig. 5E**), and even the apparent amplitude adaptation in the responses to fast-drifting gratings. The model correctly predicted some surprising aspects of measured responses, such as the different phase relations between PD and ND responses in the different cells (most cells show out-of-phase responses like Example cell 2, while a few showed in-phase responses, like Example cell 1, **Fig. 5D**). The correspondence between prediction and measurement (**Fig. 5E**) suggests that this dramatic phase difference is largely a consequence of the structure of the first-order RF of each cell. In summary, a simple model that integrates fast, excitatory and spatially offset, slow inhibitory inputs predicts T5 responses to a range of visual stimuli.

### Model analysis reveals near-optimal tuning for directionally selective responses to grating stimuli

The accuracy of our simple model in predicting grating responses encouraged us to more broadly explore the response properties of this model class in an effort to connect our work to previous thinking about insect motion detection. The responses of behaving flies and motion-sensitive neurons in flies to drifting periodic gratings have long been used to characterize the properties of the motion detectors (Buchner, 1976; Egelhaaf et al., 1989; Götz, 1968; O’Carroll et al., 1996). This work lead to the counter-intuitive finding that insect motion vision is not exclusively tuned to the velocity of a motion pattern, but rather it is more tuned to the temporal frequency (the speed divided by the spatial frequency) of the stimulus (Borst and Egelhaaf, 1989; Eckert, 1980). Correlation-type models (like Hassenstein and Reichardt (1956) or Barlow and Levick (1965)), respond to a combination of the stimulus velocity and pattern (Egelhaaf et al., 1989; Reichardt, 1987), effectively being tuned to the temporal frequency of the stimulus. Since our model was based on fitting the local flash response (**Fig. 4B**) but captured much of the response to drifting gratings (**Fig. 5**), we asked whether this type of model is also tuned to the temporal frequency of a drifting grating. To explore this model class, we made several simplifications to the model of **Fig. 4A** (see Methods; **Fig. 6A**), which slightly reduced the accuracy of the predictions, but enabled substantial clarity on the analysis of the behavior of this model. We analyzed responses to sine-wave gratings, which further simplified our analysis, and we focused on the amplitude of the periodic response (the f1 component) of the dominant contributor to responses under these conditions—the linear component of the model (numerator in **Fig. 6A**). We note that motion responses to periodic gratings have classically been analyzed at steady state (the f0 component), but here we propose that the periodic response is the more relevant factor. The classic studies focused primarily on responses of the Lobula Plate Tangential Cells (or fly behavior), which are downstream of T4 and T5 neurons. The simple spatiotemporal integration of many individual T4 and T5 neurons would result in the f1 component of the preceding layers summating and significantly contributing to the downstream steady-state response. These simplifications allowed us to describe the relationship between excitation and inhibition using only 4 (instead of 14) parameters: a time constant ratio, spatial separation factor, spread ratio, and amplitude ratio (detailed in Methods).

**Figure 6.**
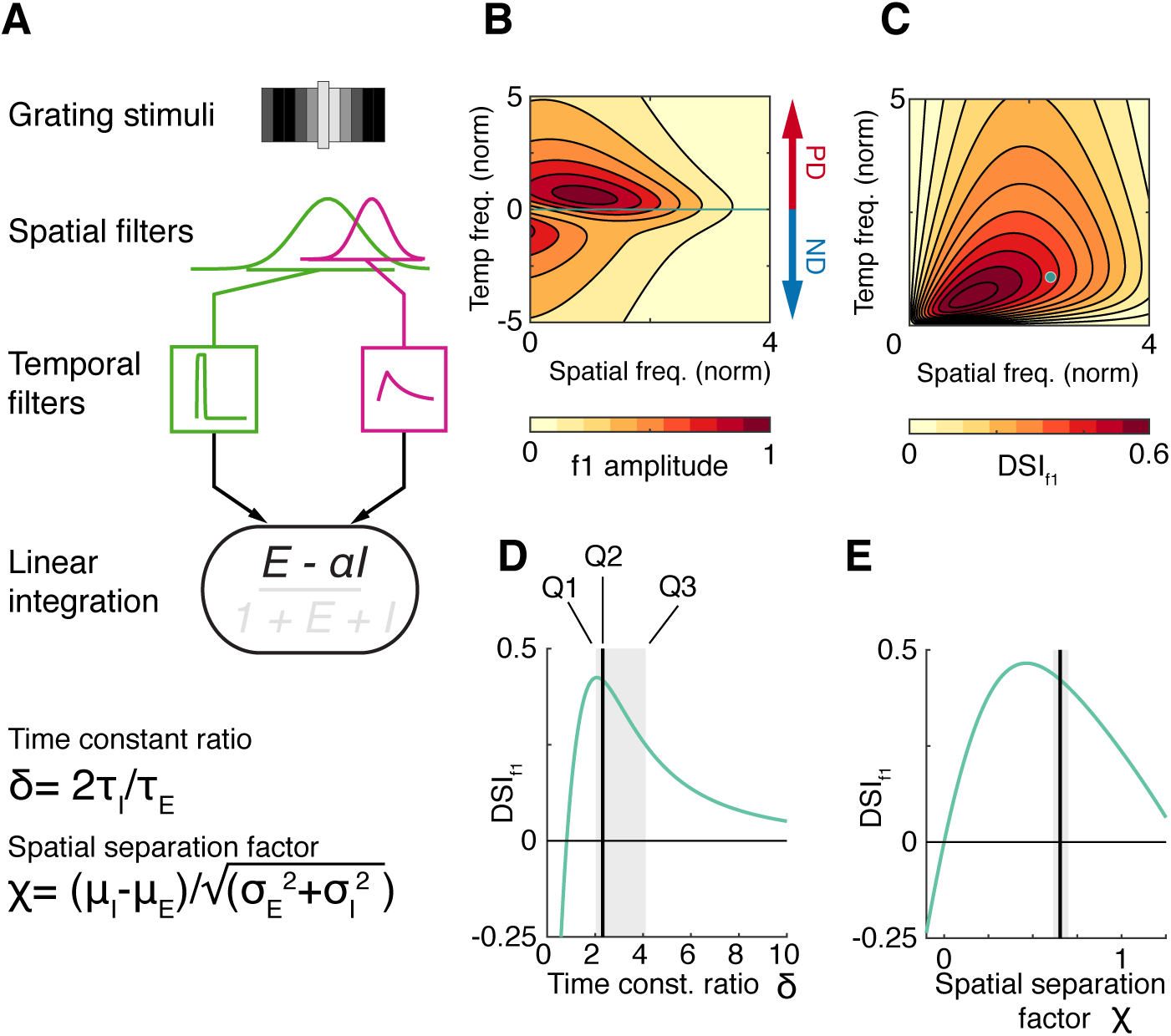
Model analysis reveals near-optimal tuning for directionally selective responses to grating stimuli. (**A**) Schematic of simplified model used to analyze responses to grating stimuli. Compared to our full model (**Fig. 4A**), the input conductances have Gaussian spatial profiles, and the output only depends on the numerator of the integration step. (**B**) Heatmap depicting the magnitude of the normalized f1 (periodic component) amplitude in response to sinusoidal grating with different spatial and temporal frequencies. The spatial and temporal frequencies are presented on a normalized (unitless) scale (see Methods). (**C**) Heatmap of the model results in **B**, but as the Direction Selectivity Index of the f1 response (DSI_f1_ = [PD_f1_ – ND_f1_]/ PD_f1_). Green dot represents our (slower) experimental grating parameters, which is used for further analysis in **D** and **E**. (**D**) The DSI_f1_ of the simplified model’s response to the grating stimulus, across values of δ, the time constant ratio (comparing the timing of the inhibitory and excitatory inputs). All other parameters kept constant, with values based on the median (of top) fit values for the compatible model (see Methods). The time constant ratio for our T5 recordings, based on the model fit is compared (median, black line, and lower and upper quartile range in gray rectangle). (**E**) Same as **D**, but for χ, the spatial separation factor (the normalized difference between the centers of the excitatory and inhibitory inputs).

We analyzed the responses of this model (**Fig. 6A**) to grating stimuli across spatial and temporal frequencies (**Fig. 6B**), using parameter values derived from our full model (see Methods). By examining the contour plot, we find that although the f1 amplitude is tuned to both spatial and temporal frequencies, the narrower distribution along the temporal frequency axis shows that it is primarily temporally tuned. The f1 amplitude has a clear peak around the same temporal frequency (∼1) for both PD and ND motion, with ND responses displaying a shallower peak that does not stretch into higher spatial frequencies. The offset peak between the PD and ND responses suggests a difference in the spatial tuning of PD and ND motion at any temporal frequency. Motion detectors of opposite directional preference are typically combined by downstream neurons after motion opponency (Mauss et al., 2015), a step that dramatically improves the accuracy of the PD computation. The DSI (based on the periodic component, referred to as DSI_f1_) implements an opponency step in our simplified model, and exhibits some speed tuning as evidenced by the diagonal tilt in the contour plot (**Fig. 6C**). While speed-tuning is not typically observed for rotational responses of flies (Tuthill et al., 2011) or motion-sensitive neurons (Eckert, 1980), we propose that a downstream cell can simply integrate opposing T4 and T5 cells to get a response with some velocity tuning, as has been observed in larger flies (Straw et al., 2006) and with behavioral experiments using translational visual motion (Creamer et al., 2018). In contrast to a previous model for generating speed tuning (Creamer et al., 2018), we do not need to invoke units with different spatial scales, since our model already features imbalanced spatial tuning for PD and ND responses arising from the spatiotemporal asymmetry between the excitatory and inhibitory inputs.

Finally, we asked how well are T5 neurons, when viewed through the lens of our simplified model, tuned to detect the direction of a drifting grating? We explored the sensitivity of our model’s directionally selective response (DSI_f1_) to variations of single parameters and based this analysis on a single grating stimulus that was matched to the stimuli we presented in our experiments (**Fig. 6C**, green dot). As our present and previous (Gruntman et al., 2018) studies have shown, DS arises from inhibition that outlasts excitation, and so we expect a strong dependence on the ratio of the inhibitory to excitatory time constants (**Fig. 6D**). Consistent with this expectation, when the ratio is close to 1, the model does not generate a directionally selective response, and the maximal directionally selective response is achieved when the inhibitory time constant is roughly twice that of excitation. For larger values of this ratio, inhibition dominates responses to any periodic stimulus, and so DS is reduced. When inhibition is faster than excitation (ratio < 1), DS reverses direction because fast inhibition interferes with integration for PD stimuli but fades quickly during integration of ND stimuli. While the equivalent computation in T5 neurons could take on any balance of temporal integration across this range, we find that the parameters fit to T5 responses sit very close to the optimal ratio with an inhibitory time constant that is approximately twice as long as the excitatory one (**Fig. 6D**, gray range). In addition to a temporal asymmetry, our model also requires a spatial offset, which in this simplified model is captured by the spatial separation factor, the normalized difference between the mean positions of the inhibitory and excitatory spatial gaussian distributions. DSI_f1_ is strongly tuned by this spatial relationship (**Fig. 6E**), with a peak response for a separation of ∼0.5. When the separation is zero (overlapping inhibition and excitation), the model is motion blind; when inhibition precedes excitation on the leading side of the RF, the preferred direction is now reversed (DSI_f1_ <0). Again, we find that the parameters fit to T5 responses reside very close to the optimal separation for this model.

## Discussion

In this study we used whole-cell recordings of T5 cells, the OFF DS neurons in *Drosophila* (**Fig. 1**) to uncover the mechanism underlying the generation of directionally selective motion responses. Using local bar flashes, we mapped the first-order receptive field of T5 neurons and revealed an asymmetric spatial structure comprised of offset excitatory and inhibitory input fields (**Fig. 2**). Using pairs of bar flashes, we mapped the responses to second-order stimuli, and found no amplifying pairwise interactions that are indicative of a PD enhancing mechanism, rather we only found evidence for ND suppression (**Fig. 3**). We modified our previous biophysical model (Gruntman et al., 2018) to better approximate single-neuron responses, and used this model to accurately predict T5 responses to static and moving bars of different widths, and even static and drifting gratings (**Fig. 4, 5**). Finally, we analyzed the behavior of our computational model and found that T5 neurons are nearly optimally tuned to compute DS (**Fig. 6**).

The first-order RF of T5 neurons, with a broad excitatory field leading a narrow trailing inhibitory field, strongly resembles the structure we uncovered for T4 neurons (**Fig. 2D**). This RF characterization agrees with recent measurements made using voltage imaging of T5 neurons (Wienecke et al., 2018), but augments them with absolute response magnitudes and time constants. The second-order response mapping was used as an explicit test for non-linear interactions that facilitate motion detection. Across a very large set of stimuli, we find a clear pattern in the responses: no evidence for an amplifying non-linearity, but abundant evidence for sublinear integration on the trailing side of the RF due to asymmetric inhibitory inputs. A recent study reported PD enhancement only occurs for larger (>6°) stimuli (Haag et al., 2017), however, even with wide bar stimuli (that nearly cover the T5 RF), we did not observe any instances of PD enhancement in the membrane potential responses (**Fig. 3D,E** and **Fig. 3 - Fig. Supp. 1**). A potential resolution to this discrepancy may be the differences in voltage versus calcium responses. When both calcium and voltage responses were imaged in T5 neurons, Wienecke et al. found amplification only in the Calcium signal (2018). Therefore, this amplification may arise in the transformation from membrane voltage to fluorescent calcium signals, however the significance of this transformation for DS computation remains an open question.

In the *Drosophila* visual system, motion responses are computed along two mostly-parallel ON and OFF pathways. The first neurons to generate directionally selective responses in their corresponding pathways, T4 and T5, have their dendrites in distinct neuropils and thus have few common inputs, and yet they share a high degree of anatomical similarity (Fischbach and Dittrich, 1989; Shinomiya et al., 2019; Strausfeld and Lee, 1991; Takemura et al., 2017; 2014). Focusing only on their inputs, both neuron types receive 4 major columnar inputs, both preferentially synapse onto neurons of their same subtype, and both receive prominent input from a single, giant GABAergic interneuron called CT1 (Shinomiya et al., 2019). However, the described anatomy of the T4 and T5 inputs combined with transcriptomic data for all major input cell types (Davis et al., 2018), suggests major differences between the 2 pathways. While T4 receives mixed inputs from columnar cholinergic, GABAergic, and Glutamatergic cells, all of the small-field inputs to T5 neurons are cholinergic. This substantial difference between the inputs of these two cells is especially surprising since the DS generation mechanism in T4 and T5 is all but identical – fast excitation with asymmetric slow inhibition on the trailing side of the RF. The most likely explanation is that T5 neurons must also receive inhibitory inputs that respond to local visual stimuli. Due to its morphology, CT1 was thought to provide a non-local inhibitory input, but a recent study raises the prospect that the local input from this neuron may function like that of other small-field inputs (Meier and Borst, 2019). A further possibility is that some fraction of the cholinergic inputs could activate an inhibitory conductance (firsy proposed in Shinomiya et al. (2014)).

While the computational mechanism is broadly common between T4 and T5 neurons, we did measure some differences between these cell types. Most notably, T5 responses showed a position-dependent rise time that was slower on the leading side than on the trailing side (**Fig. 2C**), whereas T4 neurons did not (Figure 2D of Gruntman et al. (2018)). Since T5 dendrites are retinotopically aligned with the preferred direction of motion, and since different input types synapse onto different locations of the T5 dendrite (Shinomiya et al., 2019), it is possible to connect the available functional data from these input types (Arenz et al., 2017; Serbe et al., 2016) to their input position. Indeed, Tm9 neurons, that synapse onto the distal branches of T5 dendrites (corresponding to the leading side of the RF), are also the slowest among the 4 major input types (Arenz et al., 2017). However, we found no evidence that this spatial change in rise time plays any functional role in the computation of DS. The excitatory part of the RF is ‘motion blind’ (**Fig. 3**) and when we modified our model to account for this temporal difference, we recorded no substantial improvement in model performance (results not shown). Perhaps a more important difference between T4 and T5 neurons is the sensitivity to weak stimuli.

Responses to single bar flashes in T4 were roughly equivalent to T5 responses to stimuli of twice the width (**Fig. 2D**). Haag et al. (2017) also reported a similar difference. Since this difference is already measured in response to non-moving bar flashes, it is likely a difference in the response sensitivity of upstream neurons. This difference may also suggest different detection thresholds in the ON and OFF motion pathways, which would in turn suggest different behavioral responses to small moving dark and bright objects.

In our previous work, we implemented ND suppression in a simple computational model that integrates spatially offset excitatory and inhibitory conductances in a passive, biophysical model (Gruntman et al., 2018). In this study we extended this framework to predict single neuron responses to a range of motion stimuli (**Fig. 4,5**). Importantly, we showed that a model designed only to predict single object impulse responses also accurately predicts grating responses (**Fig. 5**). This model concretely instantiates an algorithmic mechanism while also suggesting a biological implementation for the ND suppressing nonlinearity. We then used a simplified model to analyze the behavior of this model class and found that our DS mechanism is temporal frequency tuned (**Fig. 6B**). This approach, which started from fitting responses to a high-contrast static bar and ended with modelling smooth sinusoidal grating responses, may reconcile two frameworks for thinking about motion detectors: local edge motion detector with ‘featureless’ steady-state temporal frequency-tuned detectors. Temporal frequency tuning, a classic feature of correlation-based motion detection, has occasionally been seen as a sub-optimal algorithm since it cannot estimate the velocity of motion independent of the local stimulus pattern. Since PD responses and ND responses in our model are tuned to different spatial scales (**Fig. 6B**), we propose that a speed-correlated estimate can be generated downstream by a simple subtraction of outputs from detectors with opposing PDs (**Fig 6C**). Finally, we examined the sensitivity of the directionally selective estimate to individual model parameters. Remarkably, we find that models fit to T5 neurons use parameters that, when examined individually, are nearly optimal for detecting motion. This analysis provides an independent demonstration that this class of motion detector models, including the simplified version analyzed here, is a good approximation for the computation in T5 neurons.

## Acknowledgements

We thank A. Nern for providing the driver line and E. Rogers for help with fly husbandry. We are also grateful to J. E. Fitzgerald, K. D. Longden, and A. Zhao for comments on the manuscript. This project was supported by the Howard Hughes Medical Institute.

## Author Contributions

E.G. and M.B.R. designed experiments; E.G. performed experiments and analysis. E.G. and S.R. conducted the simulation study. E.G. and M.B.R. wrote the manuscript.

## Materials and Methods

### Electrophysiology

Experiments were performed on 1-2 day old female *Drosophila melanogaster* (flies were reared under 16:8 light:dark cycle at 24°C). To target T5 cells, a single genotype was used: pJFRC28-10XUAS-IVS-GFP-p10 (Pfeiffer et al., 2012) in attP2 crossed to stable split-GAL4 SS25175 (w; VT055812-AD(attP40); R47H05-DBD(attP2)) generously provided by Aljoscha Nern in Gerry Rubin’s lab (line details with expression data available from http://splitgal4.janelia.org/). Flies were briefly anesthetized on ice and transferred to a chilled vacuum holder where they were mounted, with the head tilted down, to a customized platform machined from PEEK using UV-cured glue (Loctite 3972). CAD files for the platform and vacuum holder are available upon request. To reduce brain motion the proboscis was fixed to the head with a small amount of the same glue. The posterior part of the cuticle was removed using syringe needles and fine forceps. The perineural sheath was peeled using fine forceps and, if needed, further removed with a suction pipette under the microscope. To further reduce brain motion, muscle 16 (Demerec, 1950) was removed from between the antenna.

The brain was continuously perfused with an extracellular saline containing (in mM): 103 NaCl, 3 KCl, 1.5 CaCl_2_ 2H_2_O, 4 MgCl_2_ 6H_2_O, 1 NaH_2_PO_4_ H2O, 26 NaHCO_3_, 5 N-Tris (hydroxymethyl) methyl-2-aminoethane-sulfonic acid, 10 Glucose, and 10 Trehalose (Wilson and Laurent, 2005). Osmolarity was adjusted to 275 mOsm, and saline was bubbled with 95% O_2_ / 5% CO_2_ during the experiment to reach a final pH of 7.3. Pressure-polished patch-clamp electrodes were pulled for a resistance of 9.5-10.5 MΩ and filled with an intracellular saline containing (in mM): 140 KAsp, 10 HEPES, 1.1 EGTA, 0.1 CaCl2, 4 MgATP, 0.5 NaGTP, and 5 Glutathione (Wilson and Laurent, 2005). 250μM Alexa 594 Hydrazide was added to the intracellular saline prior to each experiment, to reach a final osmolarity of 265 mOsm, with a pH of 7.3.

The mounted, dissected flies were positioned on a rigid platform mounted on an air table. Recordings were obtained from labeled T5 cell bodies under visual control using a Sutter SOM microscope with a 60X water-immersion objective. To visualize the GFP labeled cells, a monochrome, IR-sensitive CCD camera (ThorLabs 1500M-GE) was mounted to the microscope, an 850 nm LED provided oblique illumination (ThorLabs M850F2), and a 460 nm LED provided GFP excitation (Sutter TLED source). Images were acquired using Micro-Manager(Edelstein et al., 2014), to allow for automatic contrast adjustment.

All recordings were obtained from the left side of the brain. Current clamp recordings were sampled at 20KHz and low-pass filtered at 10KHz using Axon multiClamp 700B amplifier (National Instrument PCIe-7842R LX50 Multifunction RIO board) using custom LabView (2013 v.13.0.1f2; National Instruments) and MATLAB (Mathworks, Inc.) software. Shortly after breaking in, recordings were stabilized with a small injection of a hyperpolarizing current (0-3pA) setting the membrane potential to a range between -60 to -55mV (uncorrected for liquid junction potential). Occasionally, the injected current required adjustments, but these were done prior to the acquisition of the single bar flash data. To verify recording quality, current step injections were performed at the beginning of the experiment.

### Visual stimuli

The display was constructed from an updated version of the LED panels previously described (Reiser and Dickinson, 2008). The arena covered slightly more than one half of a cylinder (216° in azimuth and ∼72° in elevation) of the fly’s visual field, with the diameter of each pixel subtending an angle of (at most) 2.25° on the fly eye. Green LEDs (emission peak: 565 nm) were used, dark stimuli (off pixels) were presented on an intermediate intensity background of ∼31 cd/m^2^.

Visual stimuli were generated using custom written MATLAB code that allowed rapid generation of stimuli based on individual cell responses. In contrast to the published stimulus control system (Reiser and Dickinson, 2008), we have now implemented an FPGA-based panel display controller, using the same PCIe card (National Instrument PCIe-7842R LX50 Multifunction RIO board) that also acquired the electrophysiology data. This new control system (implemented in LabView) streams pattern data directly from PC file storage, allowing for on-line stimulus generation. Furthermore, this new control system featured high precision (10 μs) timing and logging of all events, enabling reliable alignment of electrophysiology data with visual stimuli.

To map the receptive field (RF) center of each recorded cell, three grids of flashing dark squares (on the same intermediate intensity background) were presented at increasing resolution. Each flash stimulus was presented for 200 ms. First, a 6 × 7 grid of non-overlapping 5 × 5 LEDs (∼11°×∼11°) dark squares was presented (**Fig. 1C**). If a response was detected, a denser 3 × 3 grid with 50%-overlapping 5 × 5 LEDs (∼11°×∼11°) bright and dark squares (to further verify these were T5 Cells) was presented at the estimated position of the RF center. If a recorded cell was consistently responsive to the first two mapping stimuli, a third protocol was presented to identify the RF center. A 5 × 5 grid of 3×3 LED bright squares separated by 1 pixel-shifts was presented at the estimated center of the second grid stimulus. The location of the peak response to this stimulus was used as the RF center in subsequent experiments. Once the RF center was identified, the moving bar stimulus was presented in 8 directions with 80 ms step duration (equivalent to ∼28°/s). The bar was 9 pixels in height and 1, 2, or 4 pixels in width (results in **Fig. 1D,E**). When moving in the cardinal directions, the motion spanned 9 pixels. In the diagonal directions bar motion included more steps to cover the same distance (9 steps vs. 13 steps). Once the preferred direction had been estimated, dark bar flashes were presented on the relevant axis for widths 1,2 and 4. To verify full coverage of RF, this stimulus was presented over an area larger than the original motion window (at least 13 positions; results in **Fig. 2**). In addition to these stimuli, most cells were also presented with additional stimuli following this procedure. All stimuli were presented in a pseudorandom order within stimulus blocks. All stimuli were presented 3 times, except for single bar flashes which were repeated 5 times. The inter-stimulus interval was 500ms for moving stimuli and 800ms for single bar flashes (to minimize the effect of ongoing inhibition on the responses to subsequent stimuli). Other presented stimuli were:

1. *Moving bar*. after identifying the PD-ND axis, moving bar stimuli were presented along this axis using either 40ms or 160ms steps (equivalent to 56°/sec or 14°/sec respectively). Bar height was the same for the mapping stimuli and width was either 1,2 or 4 pixels (corresponding to 2.25, 4.5 or 9°). Results in **Fig. 4**.
2. *Apparent motion*. Bar pairs were presented in 2 different configurations. Either bars were of width 2 and the delay between the first and the second bar was adjusted to maintain fixed speed (i.e. correcting the temporal delay to account for the spatial difference in positions), or bars were of width 4 and the second bar was presented directly after the first, regardless of positional difference. This second configuration was meant to elicit the strongest responses possible. Results in **Fig. 3 and Fig. 3 - Fig. Supp. 1**.
3. *Flashes of different grating phases*. Square wave gratings were of a constant spatial frequency (4 pixels OFF / 4 pixels background intensity), presented in the same window size as the moving bar stimulus (9 steps for cells with PD along a cardinal direction, 13 steps for cells with PD along the diagonals). 8 phases of the grating stimuli were presented for 40ms or 160ms. Results in **Fig. 5**.
4. *Moving grating*. Square wave grating with the same properties as above were presented with phases moving either in the forward or the reverse direction (PD or ND), and the initial phase presented being either 0 or 0.5. Results in **Fig. 5**.

### Analysis

All data analysis was performed in MATLAB using custom written code. Since the T5 baseline was typically stable, we included only trials in which the mean pre-stimulus baseline did not differ from the overall pre-stimulus mean for that group of stimuli by more than 10 mV. We also verified that the pre-stimulus mean and overall mean for that trial did not differ by more than 15 mV (or 25 mV for slow moving bars, due to their strong responses). This is the same criteria we used for our previous T4 study (Gruntman et al., 2018). Responses were later aligned to the appearance of the bar stimulus and averaged (or the appearance of the bar in the central position in case of the 8-orientation moving bar). T5 cells are expected to signal using graded synapses. Consistent with this expectation, we find that T5 recordings only occasionally feature very small, fast transients (∼1-2 mV in size) that could not be verified as spikes. Therefore, we have focused our analysis on the graded (sub-threshold) components of T5’s responses.

#### Determining PD

After presenting the cell with 1 and 2 pixel wide bars moving 8 different direction at 80 ms per step (speed which was optimal for determining directional selectivity for T4 cells, (Gruntman et al., 2018)), the preferred direction for the cell was determined by a visual estimate of the responses to determine the middle of the relatively wide range of large responding directions (see **Fig. 1**). Because stimuli were presented in 45° intervals, and the tuning of T5 neurons to direction is relatively wide, the more precise method for PD estimation that was used for T4 cells (Gruntman et al., 2018) was unnecessary.

#### DSI calculation

direction selectivity index was defined as [*R*(*PD*) − *R*(*ND*)]/*R*(*PD*), with each response defined as the 0.995 quantile (a robust estimate of the max) within the stimulus presentation window.

#### Single Position Flash Response – depolarization

responses were defined as the 0.995 quantile (a robust estimate of the max) of the response during the time between bar appearance and flash duration + 75ms. If this number did not exceed 2.5 standard deviations of the pre-stimulus baseline, the response was defined as zero. For bars of width 2 and 4 the threshold was 2.7 and 2.9 standard deviations, since the responses were stronger. Standard deviation of the baseline was determined by fitting a gaussian to the pre-stimulus baseline values for all the stimulus presentations and extracting the sigma value from the fit.

#### Single Position Flash Response – hyperpolarization

same as above only the time window used was until the end of the trial (due to slower time course for inhibition) and lower thresholds were used (1.5, 1.7, and 1.9 standard deviations, due to lower magnitude of hyperpolarization). These calculations were used for **Fig. 2C,D**.

#### Rise start/Rise time Calculation

Only presentations in which the average SPFR was detected as depolarizing were used for this calculation. Start time was defined as the time from stimulus presentation to 10% of the of the value of the maximal response for that position. Rise time was defined as the time from 10% of the response maximum to 50% of the maximum. The data in **Fig. 2C** are plotted as relative to the center (0) position, since the above values for the center position have been subtracted from all other positions for each cell separately. For each position in **Fig. 2C** at least 10 cells passed the selection criteria.

#### Apparent motion linear approximation (superposition)

Single bar flash responses were aligned to the time of the corresponding position appearance in the apparent motion stimulus. Responses were padded with zeros (since all were baselines subtracted) to extend brief single bar responses to the timescale of apparent motion. This procedure was used for the linear estimation throughout **Fig 3**.

#### Mean absolute deviation (MAD) calculation

For each stimulus presentation the mean absolute deviation was calculated between the maximum of the mean response to the stimulus, and the maximum of the individual repeats. Once a maximal response was identified for the mean response, the maximal response was found for each corresponding repeat within a 100ms window surrounding the mean peak response. We verified that there was no relationship between the maximal response magnitude and the MAD magnitude (by calculating MAD for different magnitude range) and pooled all the MAD estimates from all the different stimuli and all the cells for a global estimate. **Fig. 4C, E** and **5B, E** are using the upper quartile for a MAD estimate for all the cells. In **Fig. 4 - Fig. Supp. 1** we use the upper quartile from all the stimuli for each neuron separately.

#### Data selection

For **Fig. 5E**, only cells that showed a non-diagonal preferred direction (n=5) are presented. Since we regard the receptive field as one dimensional (PD-ND axis) and since our LED arena is limited when generating diagonal stimuli, we chose to focus only on non-diagonal PD cells instead of fitting a second dimension to the receptive field of diagonal cells.

### Statistics

To determine statistically significant differences, the one-sided, unpaired Student’s *t*-test was used for comparing groups **(Fig. 1E, 2C**, and **3D**). In **Fig. 3D**, we controlled for the false discovery rate using the Benjamini and Hochberg procedure (Benjamini and Hochberg, 1995), with q = 0.075. We noted that data were approximately normally distributed in general, but no formal test was conducted. Regression analysis for **Fig. 3E** was performed using MATLAB fit function, fitting a first-degree polynomial. No statistical methods were used to pre-determine sample sizes, however our sample sizes are similar to those reported in previous publications (Bahl et al., 2015; Turner-Evans et al., 2017; Tuthill et al., 2014). Data collection and analysis could not be performed blind to the conditions of the experiments.

### Data Plotting Conventions

All boxplots presented were plotted with these conventions: box represents upper and lower quartile range, line represents median, whiskers were omitted, and individual data points are overlaid on the box.

### T5 neuron model

#### Model dynamics

We modeled the dynamics of T5 somatic membrane potential, *V*(*t*), as a single-compartment conductance-based neuron

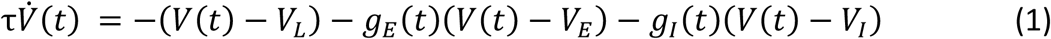

where τ denotes the integration time constant of the model neuron and *V*_*E*_, *V*_*I*_, *V*_*L*_ denote, respectively, the excitatory, inhibitory, and leak reversal potentials. The dynamics of the excitatory and inhibitory conductances (*g*_*k*_(*t*); *k* = *E, I*), measured in units of leak conductance, are described by two first-order linear filters in series

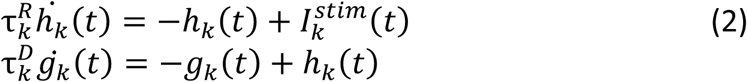

where 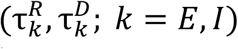 denote the rise and decay time constants of the conductances, and 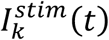 is determined by the spatial receptive fields of the conductances and the spatiotemporal profile of the visual stimulus.

#### Model inputs

The input to the conductance equations

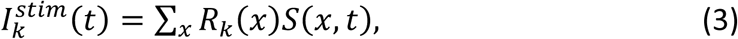

is determined by the spatiotemporal profile of the visual stimulus *S*(*x, t*) and the spatial receptive field of the conductance (*R*_*k*_(*x*); *k* = *E, I*). The integer index *x* runs over locations along the ND-PD axis of a cell. This spatial discretization is dictated by the smallest size of the bars used in the experiment (width 1). The arbitrary reference *x* = 0 denotes the empirically determined location in the cell receptive field where a flashed bar elicited the strongest depolarization. The stimulus *S*(*x, t*) can assume two possible values, 0 or 1, denoting respectively the absence or presence of an OFF bar of width 1 at the corresponding spatial location and time. We are not explicitly considering the height of the bar, hence our model stimuli replicate the one-dimensional equivalent of the stimuli used in the experiments.

#### Receptive fields

A preliminary comparison between model and T5 responses indicated that the excitatory receptive field is approximately symmetric, while the inhibitory receptive field has a long tail toward the leading (excitatory) side (data not shown). Thus, we modeled the spatial receptive field of the excitatory conductance as a Gaussian profile

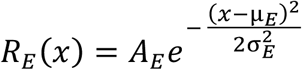

centered at location μ_*E*_, with amplitude *A*_*E*_ and width σ_*E*_. The spatial receptive field of the inhibitory conductance was described by the asymmetric profile

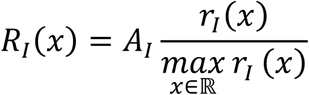

Where *A*_*I*_ is the maximal conductance and

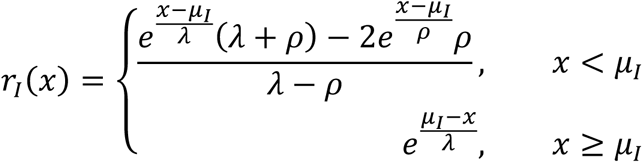

The location of the inhibitory receptive field along the ND-PD axis is controlled by μ_*I*_, the width is determined by both ρ and λ, while the amount of asymmetry is determined by ρ.

#### Model optimization

We numerically integrated the model dynamics with a 4^th^-order Runge-Kutta integration scheme. We obtained model parameters by numerically minimizing the squared error between the membrane potential dynamics of the model and the measured membrane potential of T5 cells in response to flashed bars of width 2. A constrained minimization procedure was performed 1000 times for each cell, starting from random uniform initialization of the parameter values within specified bounds (Supplementary Table 1). Predictions for the remaining sets of stimuli from the top 1% of models (10/1000, based only on the magnitude of the error for the width 2 bar flash stimuli) were then compared with the measured responses (**Figs. 4,5**). All simulations were performed in Matlab. Code and data will be made available online.

### Simplified T5 neuron model – continuous gratings

For the modeling results related to drifting gratings presented in **Fig. 6**, we considered a simplified version of the model described above, making several simplifying assumptions: (i) the time constant of integration of the neuron model is negligible compared to the other time scales in the model, (ii) the model response to drifting gratings is approximately linear (Wienecke et al., 2018), and (iii) spatial and temporal receptive field were further simplified to allow an analytical solution to the model. We describe these three steps below. We used the resulting simplified model to explore the response properties of this model class to a broader stimulus space beyond the stimulus set used in the electrophysiological experiments.

#### (i) Model simplification

The constrained optimization procedure consistently produced sets of model parameters where the time constant of neuronal integration τ was close to the lower bound of the imposed constraints (∼1ms). Given the negligible time constant of integration for the T5 model neurons, the membrane potential dynamics can be well described by a model with instantaneous neuronal integration

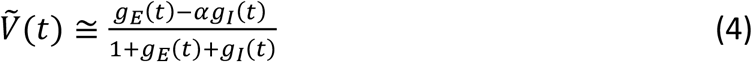

where 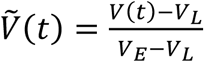 and 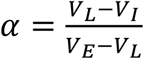 (see e.g. Gruntman et al, 2018).

#### (ii) Analysis of the model response to drifting gratings

We now consider the idealized scenario of a spatially and temporally continuous drifting sinusoidal grating stimulus, with temporal frequency ω and spatial frequency *k*

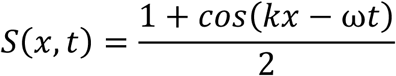

Note the slight abuse of notation, as temporal frequency is 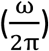 and spatial frequency 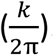. Given the linear relationship between the stimulus and the excitatory and inhibitory conductances (Eqs. 2,3), each conductance can be described by the amplitude of its 0-th Fourier coefficient 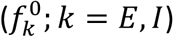, the 1^st^ Fourier coefficient 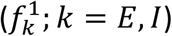, and a phase (Ψ_*k*_; *k* = *E, I*). The resulting time course of the membrane potential (Eq. 4) becomes

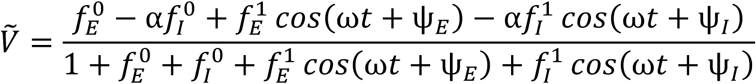

The two oscillatory terms in the numerator (N) and denominator (D) can be further combined to yield new terms described by the amplitude of the resulting sinusoidal oscillations 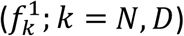 and their phases (*Ψ*_*k*_; *k* = *N, D*)

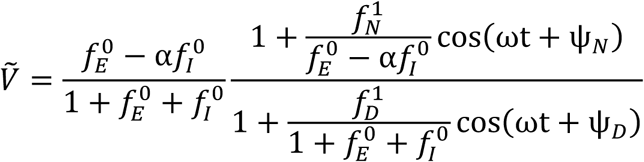

Hence, in order to estimate the oscillation amplitude of the membrane potential (up to scaling factors that are irrelevant when considering ratios, such as DSI), we need to compute the 1^st^ Fourier component of the function

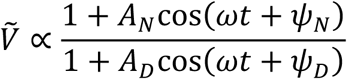

where 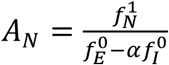 and 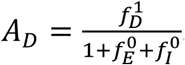. This Fourier component can be computed analytically

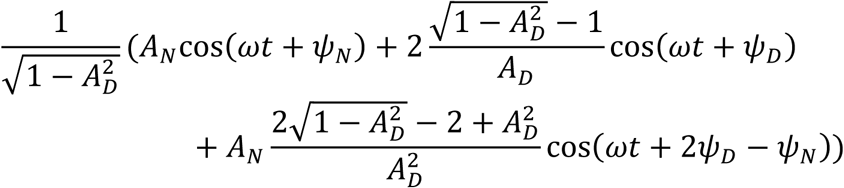

We know that, by definition, the denominator in Eq. 4 can never become negative, hence the amplitude of the oscillatory term in the denominator needs to satisfy *A*_*D*_ < 1. Moreover, given the observed hyperpolarizing T5 responses to grating stimuli (**Fig. 5**), the amplitude of the oscillatory term in the numerator needs to be sufficiently large to induce a change of sign in the numerator, *A*_*N*_ ≳ 1. Under these conditions, the dominant contribution to the oscillation amplitude of the membrane potential comes from the first term, *A*_*N*_cos(*ωt* + *Ψ*_*N*_). Therefore, it is sufficient to estimate the first Fourier component of the numerator, 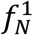, which in turn can be estimated from the first Fourier component of the excitatory and inhibitory conductances.

#### (iii) Analysis of the linear model response to drifting gratings

The optimization procedure resulted in models where the rise time of the excitatory conductance 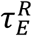 was close to the lower bound of the imposed constraints (∼1ms). The optimization results also revealed that the rise and decay times of the inhibitory conductance were tightly correlated, such that their sum is approximately preserved. For simplicity, we only considered the regime in which these two time constants (τ_*I*_) are identical. We further simplify the model by assuming a Gaussian spatial receptive field for the inhibitory conductance,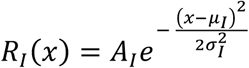.

With these additional simplifications, the first Fourier coefficient of the numerator in Eq. 4, 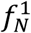, becomes

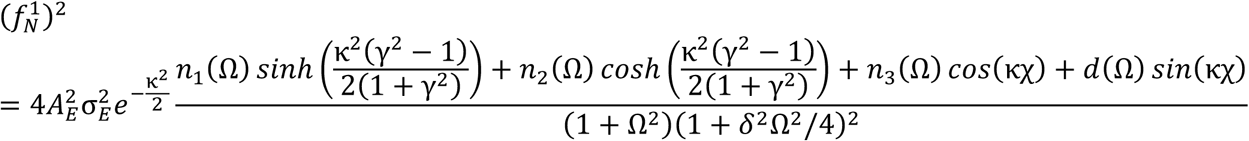

Where

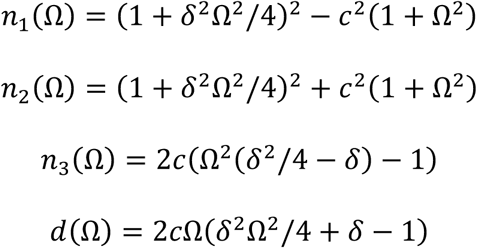

In these equations (i) *δ* denote the ratios between inhibitory and the excitatory time constant 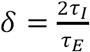 (ii) 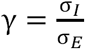 is the ratio between the size of the receptive fields (iii) 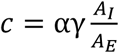 measures the relative strength between the inhibitory and the excitatory drive (iv) 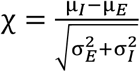 measures the separation between the excitatory and inhibitory receptive fields (v) 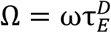 is the normalized temporal frequency, expressed in units of the inverse excitatory time constant, and (vi) 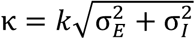 is the normalized spatial frequency, expressed in units of the inverse overall spread of the two receptive fields.

In order to estimate the normalized parameter values, we first optimized a new model of T5 responses that is compatible with the assumptions of the simplified model described in this section: (i) The spatial receptive field for the inhibitory conductance was Gaussian (ii) We only considered optimization results where the neuronal integration time was small (<5ms) (iii) we verified that the sum of the rise and decay time for the inhibitory conductance was approximately constant across optimization results. Typical values for the normalized parameters from the optimized models are: *δ* = 1, *γ* = 1.1, *c* = 0.74, *χ* = 0.65. The quantity 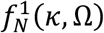, with peak normalized to 1, is depicted in **Fig. 6B**. Note that the only directionally selective term in 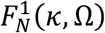 is *d*(Ω) *sin*(κχ).

Finally, we estimate the DSI of the f1 component as 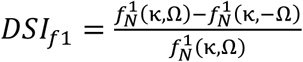 (see **Fig. 6C**).

**Supplementary Table 1:** Model parameters, related to Fig. 4 and 5.

| Parameter | Description | Units | Bounds |
| --- | --- | --- | --- |
| $\tau_E^R$ | Rise time of E conductance | ms | 1 - 400 |
| $\tau_E^D$ | Decay time of E conductance | ms | 1 – 400 |
| $A_E$ | Conductance amplitude (E) | unitless | 0 – 10 |
| $\mu_E$ | Receptive field location (E) | unitless | -5 – 5 |
| $\sigma_E$ | Width of conductance spatial profile (E) | unitless | 0 – 10 |
| $\tau_I^R$ | Rise time of I conductance | ms | 1 - 400 |
| $\tau_I^D$ | Decay time of I conductance | ms | 1 - 400 |
| $A_I$ | Conductance amplitude (I) | unitless | 0 – 30 |
| $\mu_I$ | Receptive field location (I) | unitless | 0 – 200 |
| $\lambda$ | Width of conductance spatial profile (I) | unitless | 0 - 10 |
| $\rho$ | Asymmetry of conductance spatial profile (I) | unitless | 0 - ∞ |
| $\tau$ | Integration time of neuron | ms | 1 - 400 |
| $V_E$ | Reversal potential (E) | mV | 0, Fixed |
| $V_I$ | Reversal potential (I) | mV | -80 – -65 |
| $V_L$ | Resting potential | mV | -65, Fixed |

---

**Figure 3 – Figure supplement 1.**
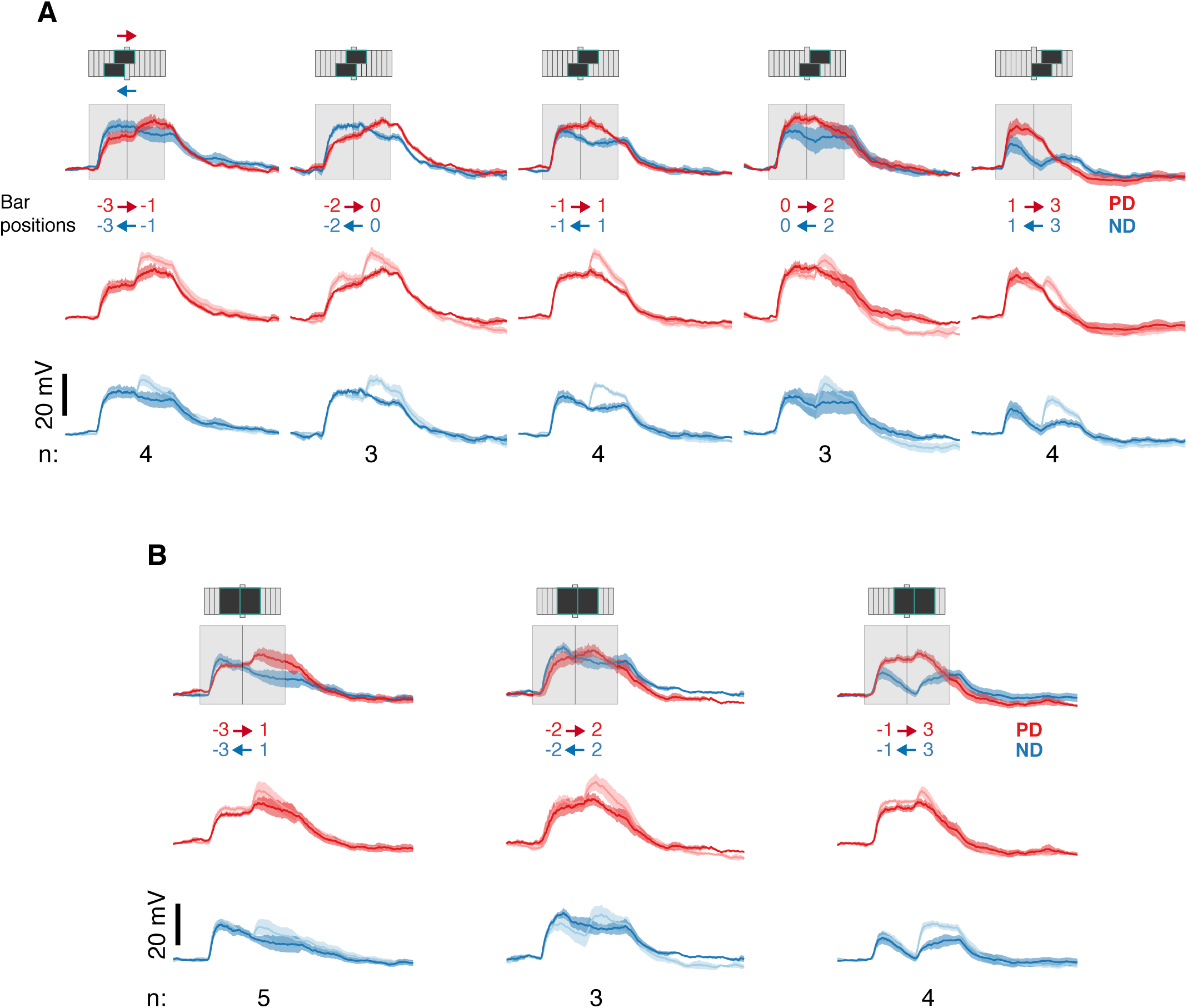
Apparent motion responses show evidence only for ND suppression even for larger stimuli. (**A**) Same as **Fig. 3B** for 4-pixel (9°) wide apparent motion stimuli. Stimulus schematic depicts positional information only and are presented in a staggered manner to illustrate overlapping positions for these pairwise combinations. Numbers below each column indicate the number of cells for that pairwise comparison. (**B**) Same as **A**, with non-overlapping positions. Summary of these pairwise combination is shown in **Fig. 3D**. Every combination shows a clear sub-linearity in response to the second bar’s appearance (measured response is always close to or smaller than the summed response in the lighter color).

**Figure 4 – Figure Supplement 1.**
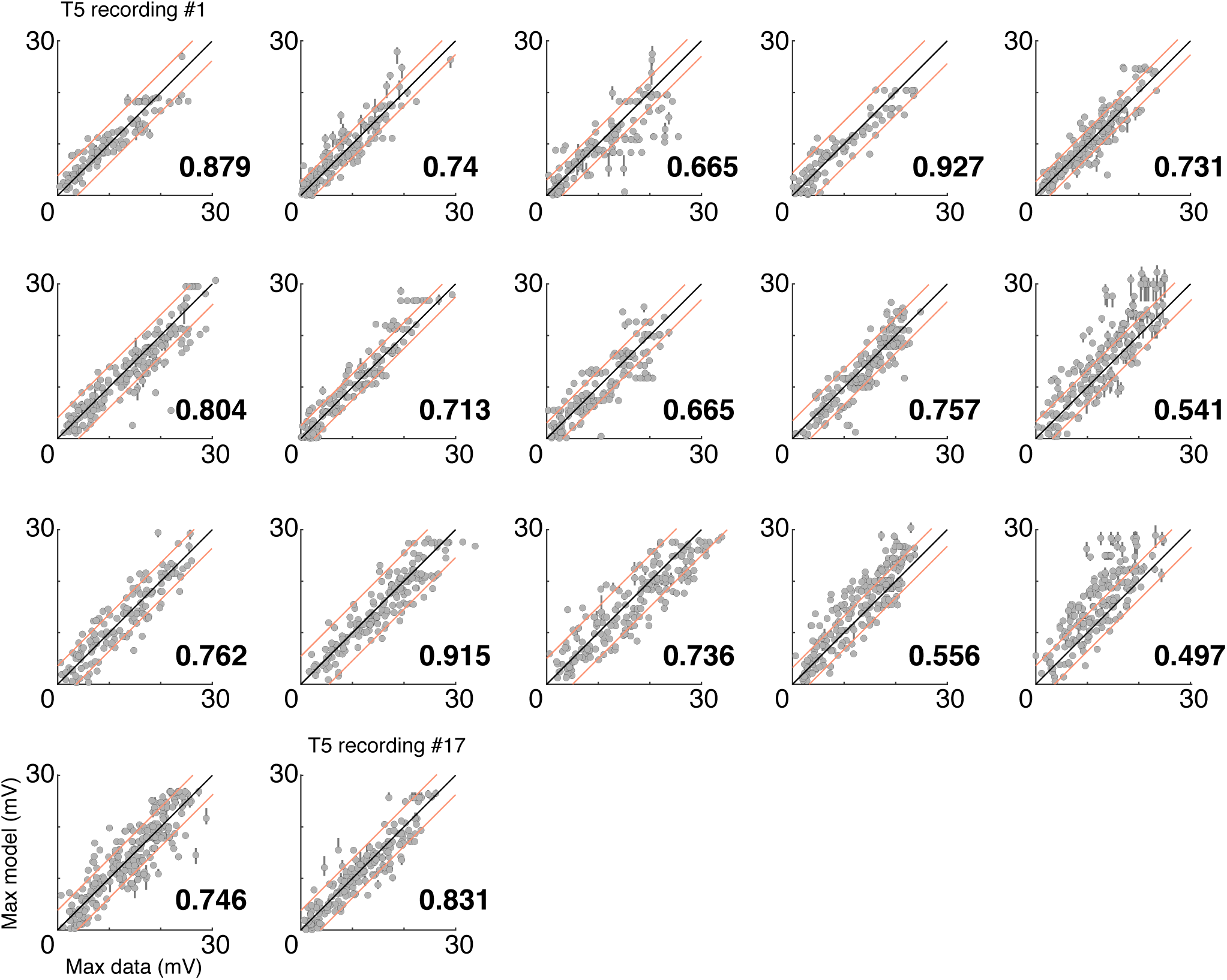
Reliability of model predictions across cells. Each subplot shows peak measured responses compared to the peak model prediction responses for all the stimuli recorded for an individual cell. Plotting conventions are as in **Fig. 4C**. The bolded value on the lower right of each plot is the proportion of model responses falling within the ‘MAD bounds’ (a measure of dispersion bases on the Mean Absolute Deviation of measured responses to the same stimulus).

